# Bending-torsional elasticity and energetics of the plus-end microtubule tip

**DOI:** 10.1101/2021.08.12.456048

**Authors:** Maxim Igaev, Helmut Grubmüller

## Abstract

Microtubules (MTs), mesoscopic cellular filaments, grow primarily by the addition of GTP-bound tubulin dimers at their dynamic flaring plus-end tips. They operate as chemomechanical energy transducers with stochastic transitions to an astounding shortening motion upon hydrolyzing GTP to GDP. Time-resolved dynamics of the MT tip – a key determinant of this behavior – as a function of nucleotide state, internal lattice strain, and stabilizing lateral interactions have not been fully understood. Here, we use atomistic simulations to study the spontaneous relaxation of complete GTP-MT and GDP-MT tip models from unfavorable straight to relaxed splayed conformations and to comprehensively characterize the elasticity of MT tips. Our simulations reveal the dominance of viscoelastic dynamics of MT protofilaments during the relaxation process, driven by the stored bending-torsional strain and counterbalanced by the inter-protofilament interactions. We show that the post-hydrolysis MT tip is exposed to higher activation energy barriers for straight lattice formation, which translates into its inability to elongate. Our study provides an ‘information ratchet’ mechanism for the elastic energy conversion and release by MT tips and offers new insights into the mechanoenzymatics of MTs.

## Introduction

Microtubules (MTs) span the intracellular space, define the position of the nucleus, build the spindle needed to segregate genetic material, and define the shape of axons, cilia as well as transport networks [1]. They are dynamic non-covalent polymers formed by tubulin dimers that hydrolyze GTP to GDP upon self-assembly. The polymerization dynamics and mechanical properties of MTs — which determine both network architecture and directional force generation — are central to cell physiology. MTs are structurally stiff due to the large and hollow cross-sections, but highly dynamic at their ‘plus-end’ tips, while the ‘minus-end’ tip dynamics are usually suppressed by MT nucleation complexes [2]. During growth, the plus-end tips stochastically transition to periods of rapid shortening (referred to as *catastrophe* events), depending on polymerization conditions, the shape raggedness of MT ends, the content and distribution of GTP-tubulin in the lattice, and occasionally, on post-translational modifications and MT-associated proteins [3,4,5,6]. This ‘power stroke’ provides the directional force that drives, particularly, the segregation of chromosomes during mitosis [7,8,9]. The elastic energy released by MT protofilaments (PFs) curling outwards is transmitted mechanically to kinetochore-MT ring complexes [10,11,12,13], which translates into directional and concerted chromosome movement.

With its well-separated modes of operation – chemo-mechanical energy conversion, storage, and release – this protein engine is an excellent object for studies on nanomechanics. The ultimate goal is to understand the functioning of this astounding out-of-equilibrium machine at the atomic scale. The kinetics of MT assembly and catastrophe have been fairly well characterized [14,15,16,17,18] compared to the MT tip structure, the dynamics of which are hard to time resolve at high resolutions with modern techniques and despite its critical role in setting up these kinetic rules. Although an early cryoelectron microscopy (cryo-EM) study [19] found that MT tips are rather blunt when growing and flared when shrinking, further studies reported a much wider range of shapes for growing MT tips [20,21,22,23,24]. Meanwhile, a model of consensus is emerging in which MTs have ragged and splayed tips with PFs curling outwards from the lumen, irrespectively of their polymerization state [25,26,27].

Despite the wealth of available structural data, the mechanistic details of this energy conversion and transduction are still not fully understood. It is known, from optical tweezers assays [7,8,28] and force-velocity measurements [29,30], that disassembling MTs (most likely, almost exclusively GDP-MTs) are capable of developing large pulling forces in the range of 30 – 65 *pN* in the presence of load-bearing attachment to kinetochores, while growing MTs (most likely, densely capped by GTP-tubulin) can generate pushing forces up to 3 – 4 *pN* through incorporation of tubulin dimers (Fig. 1*A*). What still remains enigmatic is, however, why these opposing behaviors exist if growing and shrinking MTs apparently have very similar tip structures and what role the flared tip morphology plays in the mechanism of MT catastrophe. The interplay of intrinsic PF strain and stabilizing lateral interactions is complex, and very few is known about the transient dynamics of the MT tip along its internal degrees of freedom that might be strongly affected by GTP hydrolysis but have not been resolved previously. A clear mechanochemical picture that fairly describes all these phenomena is not yet available.

**Figure 1:**
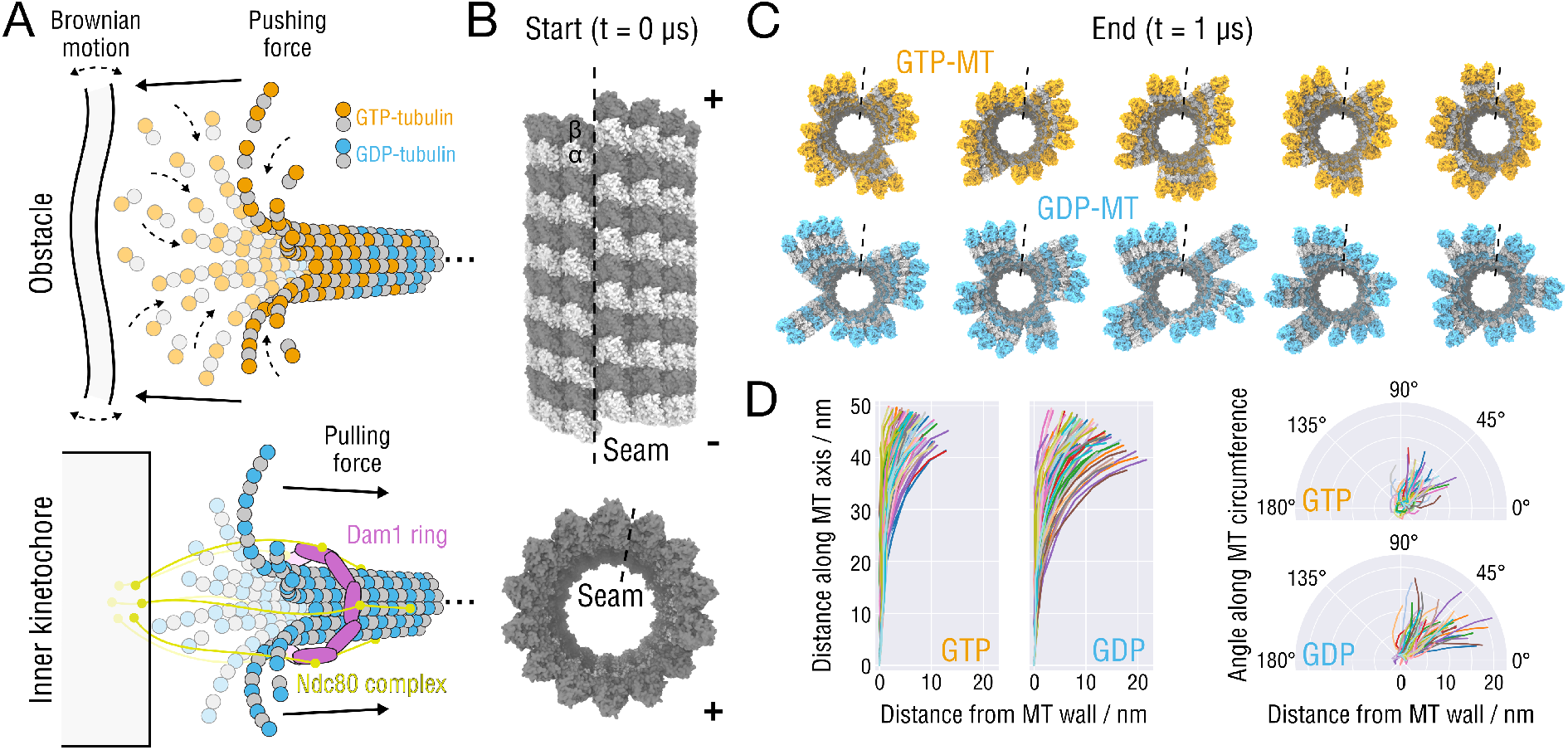
Splaying relaxation of the MT plus-end tip. **(A)** Biochemical basis of force generation by polymerizing and depolymerizing MTs (adapted from [10,8,13,25]). (*Top*) A MT growing against an intracellular ‘obstacle’, e.g., lipid membranes, organelles or other protein complexes. Stochastic fluctuations allow for incorporation of tubulin dimers into the lattice, leading to a displacement or deformation of the ‘obstacle’ (assuming the minus-end is fixed). (*Bottom*) Schematic showing the attachment of the MT tip to the Ndc80-Dam1 kinetochore complex that is thought to slide toward the minusend using the mechanical force transmitted from peeling PFs during MT depolymerization (assuming the minus-end is fixed). **(B)** Side and top views of the starting MT tip structure. Atomistic structures are shown in surface representation. The lowest row of tubulin monomers at the minus-end were restrained during the simulations. The position of the seam is indicated with a dashed line. **(C)** Top view of the GTP-(orange) and GDP-MT (blue) tip structures after 1 *μ*s of simulation. The seams are indicated with dashed lines. **(D)** Traces of the PFs aligned with respect to the minus-end monomer and projected onto the radial (left) and axial (right) planes. Random colors were assigned for clarity.

To answer these questions, we carried out atomistic molecular dynamics (MD) simulations of the *complete* MT plus-end tip both in GTP- and in GDP-state starting from an initially straightened MT structure (all PFs are straight) and analyzed the subsequent splaying of individual PFs driven by their outward curling. We assumed that this relaxation dynamics is directly related to the twist-bending stiffness of individual PFs and inversely related to the lateral interaction free energies. Quantifying the elementary intra- and inter-PF contributions to the overall thermodynamics and kinetics of MT tip splaying allowed us to infer the basic energetic criteria for the MT plus-end tip stability and to propose a mechanism of MT catastrophe that does not involve differences in the shapes of growing and shrinking MT tips.

## Results and Discussion

### Relaxation dynamics and heterogeneity of MT plus-end tips

At the MT tip, the elastic energy stored by straightened PFs is only insufficiently balanced by the total energy of lateral dimer-dimer interactions, which results in MT structures featuring irregularly splayed, curved PFs [25,26,27]. Recent atomistic MD simulation have predicted substantial differences in the bending rigidity of GTP- and GDP-bound free dimers, single PFs, and tubulin octamers [31,32,33,34,35,36]. Concomitantly, it has been proposed that the formation/breakage of lateral dimer-dimer contacts might also be nucleotide-dependent [37,38,39]. Hence, we first asked if the nucleotide state affects the overall dynamics of *complete* MT tips, where all these factors come together and can be assessed simultaneously.

To this end, we constructed two initial atomistic models of GTP- and GDP-MT tips by consecutively stacking three identical MT segments, each comprising two layers of dimers and each optimized against the high-resolution cryo-EM densities of GMPCPP- or GDP-MTs [40,41,35] (see Fig. 1*B* and Methods for a detailed simulation protocol). Immobilization of the minus-end tip was modelled by harmonically restraining the lowest row of tubulin monomers to their initial positions. Relaxation of the pre-equilibrated tip models was then quantified in five independent 1-*μ*s simulations for each nucleotide state (Fig. 1*C*; Movies S1-S20 visualizing the relaxation process). The final tip structures after 1 *μ*s of relaxation qualitatively resemble experimental observations [25,26,27]. Overall, it is seen that the initial straight configuration is highly unstable. Notably, the relaxation is somewhat stochastic in that it does not follow the same ‘pathway’ in every simulation. Rather, the MT tips crack open at different lateral interfaces, with none of them being unique, except for the seam that splays faster and more strongly than other interfaces in all performed simulations (discussed in later sections).

To compare the GTP and GDP tips directly, we aligned the PFs of the final tip structures with respect to the first minus-end monomer and projected their traces onto the radial and axial plains (Fig. 1*D*; calculation of traces is explained in Methods). Our analysis shows a stronger tendency of the simulated GDP-MT tips to splay and a stronger tendency of their PFs to bend radially. Here, the obtained projections do not fully reflect the equilibrium distribution of individual PF curvatures because (a) our MT tip models are – by construction – too large to converge within 1 *μ*s (root mean square deviations (RMSDs) from the starting conformations are shown in Fig. S1), and (b) most PFs are still stabilized by lateral neighbors.

We further observed that all of the simulated MT tips additionally accumulate a twist when looked at from the plus-end perspective (Fig. 1*C,D* and Movies S1-S20). Closer inspection of the simulation trajectories showed that this motion is strictly clockwise and caused by the bent PFs twisting similarly clockwise about their centerlines. Crystallographic data [42,43,44] and previous MD studies [31,33,39] show that such torsional motions are an intrinsic property of tubulin dimers and PFs. This reflects the structurally complex nature of the bending-torsional coupling at the MT plus-end.

### Essential dynamics and energetics of PF fluctuations

The complex bending-torsional coupling and the clockwise lattice twist of the individual PFs at the MT tip, particularly pronounced in GDP-MTs, suggests that the usually employed single angular variables describing the nucleotide-dependent radial bending angles between adjacent tubulins [45,46,47,27,36] may be insufficient to fully explain the conformational dynamics of PFs. We therefore asked, what are the essential collective variables that quantitatively describe bending-torsional PF fluctuations in our atomistic simulations and exactly how are these affected by the nucleotide state?

To investigate the bending-torsional PF dynamics in more detail, we carried out additional simulations of isolated minus-end-fixed PFs in both nucleotide states and analyzed their equilibrium dynamics in more detail. Straight PFs comprising three longitudinally connected tubulins were extracted from the complete MT lattices (see Methods), and their fluctuations were then quantified in a set of 20 independent 4-μs simulations for each nucleotide state. Principal component analysis (PCA; see Methods) was performed on the backbone atoms of the PF structures aligned with respect to the minus-end monomer to capture their collective motion (Fig. 2*A*; Fig. S2 additionally shows the equilibrium distribution of PF traces projected onto the radial and axial planes).

**Figure 2:**
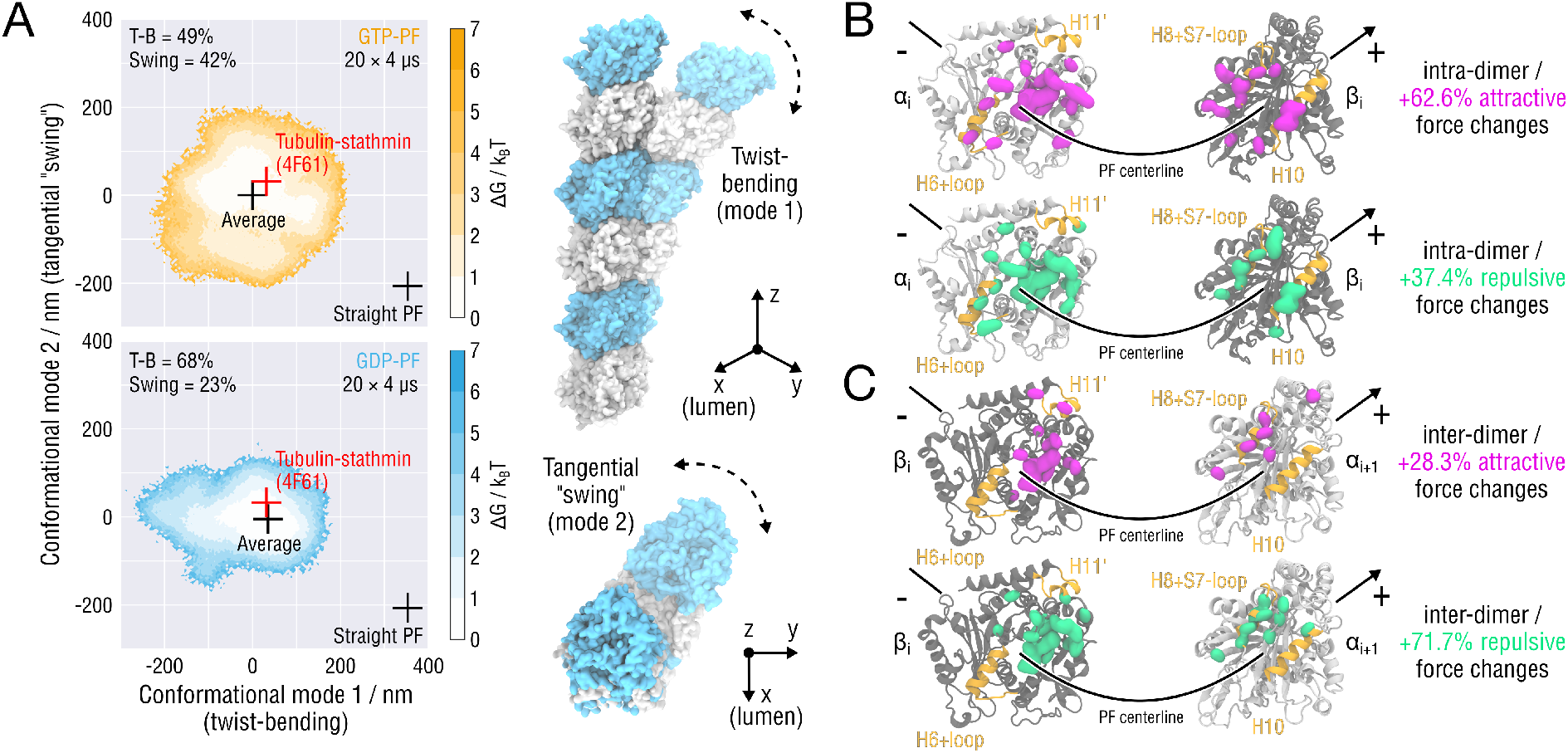
Essential dynamics of PF fluctuations and structural determinants of PF twist-bending. **(A)** Free energy landscapes of the minus-end-fixed PF conformations obtained by 20×4-*μ*s all-atom MD simulations for each nucleotide state (see Methods). Color coding as in Fig. 1. Black crosses denote the straight and relaxed (average) PF conformations. The free energy minimum corresponds to the average PF conformation. Red crosses indicate the conformation of the experimental structure of the PF oligomer co-crystallized with the stathmin protein (PDB ID: 4F61 [44]). Also shown schematically are PF rearrangements along modes 1 and 2 (see also Movies S21-S24). **(B)** Intra-dimer (top) and **(C)** inter-dimer (bottom) contact clusters and pairwise interaction forces obtained by the Force Distribution Analysis (FDA). Relative changes are shown reflecting shifts in the pairwise contact forces after GTP hydrolysis. Amino acid residues contributing to negative shifts (more attractive forces) are marked with magenta blobs, whereas those contributing to positive shifts (more repulsive forces) are marked with green blobs. A lower limit cutoff of 50 pN for both negative and positive force changes was applied to highlight high-force clusters. *α*- and *β*-tubulins are shown as silver and gray ribbons, respectively. Highlighted in yellow are residues involved in the H6 (+ loop), H8 (+ loop), H10, H11’ helices.

Two major conclusions can be drawn from this analysis. First, more than 90% of the equilibrium fluctuation variance can be explained by only two (principal) modes of motion. Mode 1 represents the bending-torsional PF motion observed in the large-scale MT tip simulations (Movies S21-S22). Mode 2 describes a tangential motion of the PF perpendicular to the radial plane (here, referred to as *tangential swing*; Movies S23-S24). To additionally verify if the minimum free energy conformations seen in our simulations agree with PF conformations observed experimentally, we also projected the crystal structure of the tubulin-stathmin oligomer complex (PDB ID: 4F61 [44]) onto our calculated free energy profiles (Fig. 2*A*, red crosses; see Methods). Importantly, bending of the PFs and twisting about their centerlines do not come out as separate eigenmodes in the PCA, but are highly correlated, which strongly suggests that using only a single bending angle describing the local radial PF curvature may miss a large part of the actual conformational dynamics of individual PFs.

Second, the conformational dynamics of PFs depends strongly on the nucleotide state. According to the PCA results, the conformational flexibility of GDP-PFs is reduced by 37% compared to that of GTP-PFs (see Methods). In order to estimate the free energy stored in straight GTP- and GDP-PFs, we fit 2-dimensional harmonic functions to the free energy landscapes in Fig. 2*A*, which yielded Δ*G*_0_ = 16.1 ± 0.1 k_B_T and Δ*G*_0_ = 25.7 ± 0.2 k_B_T, respectively. We emphasize that these estimates rest on the harmonic approximation, not on direct sampling. Further, whereas for the GTP-PFs both bending-torsional (mode 1) and tangential swing (mode 2) components explain almost equal fractions of the conformational flexibility (49% and 42%, respectively), this distribution is shifted substantially to 68% and 23% for the GDP-PFs, respectively. Neglecting the remaining variance unexplained by modes 1 and 2 (less than 10%), the decomposition provided by the PCA suggests an allosteric mechanism that couples the PF bending-torsional dynamics to the change of the nucleotide state. More specifically, the rate and the exact modality (*i.e*., stronger or weaker mode 2) would be largely changed upon GTP hydrolysis.

To the best of our knowledge, the tangential swing component has not been reported in previous MD studies [48,49,33,39], most likely, due to shorter simulation times. Speculating about its role in the plus-end tip dynamics, we propose that lower tangential stiffness might facilitate the stochastic formation of lateral PF-PF contacts through spontaneous collisions. The role of the lateral contacts in these processes will be elucidated in later sections.

### Structural determinants of the intra- and inter-dimer coupling in PFs

As the mechanical coupling along the PF shaft is a key determinant of the overall bending-torsional elasticity, we analyzed the longitudinal interactions both within and between the tubulin dimers in more detail. To this end, we identified pairs of residues that contribute the most to the intra- and inter-dimer interactions in our equilibrium PF simulations, thus forming a network that stabilizes the PF and modulates its essential conformational dynamics shown in (Fig. 2*A*). Here, we first used the time-resolved Force Distribution Analysis (FDA) [50,51] to follow the dynamics of all internal forces within the PFs in GTP- and GDP-state (see Methods). We then averaged the obtained forces and filtered out those force contributions that were unrelated to the intra- and inter-dimer contacts. By convention, negative (positive) pairwise forces are attractive (repulsive) forces; and hence, negative (positive) changes in the pairwise forces denote stronger (weaker) intra- and inter-dimer interactions.

Figure S3 shows the identified contact clusters for each nucleotide state. Two main regions exhibit strong (absolute values larger than 50 pN) pairwise residue forces. The first one is located mainly in the vicinity of the hydrophobic contact (‘anchor point’ [52]) between the H8 helix plus the accompanying H8-S7 loop of the top monomer and the H11’ helix of the bottom monomer, as well as around the nucleotide binding pocket. In this region, not all of the interactions are clustered near the putative ‘outside’ of the MT surface; some of them reach out to the putative PF-PF contact zones, which partially explains the non-radial nature of PF bending. The second region is located near the putative ‘inside’ of the MT surface. Here, the portion of the highly conserved H10 helix of the top monomer that is closer to the MT lumen interacts with the H6 helix plus the adjacent loop of the bottom monomer. Interestingly, this contact cluster is only present at the intra-dimer interface, which might explain the higher PF flexibility at the inter-dimer interfaces, as reported previously [33].

Figure 2*B,C* shows changes in the average pairwise residue forces after GTP hydrolysis. Contrary to our initial expectation that GTP hydrolysis would only affect the force distribution at the interdimer interface (indeed, the negatively charged *γ*-phosphate and Mg^2+^ are removed), we found that post-hydrolysis changes in the PF structure have opposite effects at the intra- and inter-dimer interfaces. First, the overall rearrangements near the nucleotide binding pocket result in a cumulative weakening of the inter-dimer interface: +71.7% of repulsive pairwise forces *vs*. only +28.3% of attractive pairwise forces (Fig. 2*C*). This finding correlates well with a recent MD study showing that GTP hydrolysis results in a ~4 k_B_T longitudinal bond weakening [36] as well as with our previous finding that the longitudinal bonds in GDP-PFs are more fragile when subject to mechanical stretching [35]. We note, however, that the FDA provides only the enthalpic contribution to the longitudinal stability because the entropic penalty for eliminating solvent from tubulin’s surface is not taken into account. Second, inspection of the average pairwise force changes at the intra-dimer interface points to a cumulative strengthening: +62.6% of attractive pairwise forces *vs*.+37.4% of repulsive pairwise forces (Fig. 2B). This stabilization arises mainly from rearrangements near the nucleotide binding pocket as well as from altered interactions between the H10 and H6 helices. Because the hydrolysis reaction does not happen at the intra-dimer interface [53], we speculate that the conformational changes triggered by GTP hydrolysis (lattice ‘compaction’ around the longitudinal inter-dimer interface [54,52,40,35]) propagate upstream, allosterically causing additional intra-dimer tensions.

### The role of lateral contacts in MT lattice integrity

Relaxed PFs tend to adopt curved-twisted conformations due to the heterogeneous interaction profiles at their intra- and inter-dimer interfaces (Fig. 2*A,B*). However, straight PFs need to be stabilized via specific interlocking of *α*- and *β*-tubulin monomers (Fig. 1*B-D*). This draws attention to the lateral contacts, the exact role of which in the MT stability is still controversially debated [55]. With two notable exceptions [38,39], simulation studies have proposed that the stability of lateral PF-PF contacts is unaffected by the nucleotide state [32,35,33,34,36]. However, only in one of these studies [34], the lateral binding free energy was computed explicitly using enhanced sampling techniques, though with relatively moderate sampling times. Furthermore, the available MD simulation evidence does not account for and cannot rule out that also kinetic aspects of the lateral PF-PF association play an active role in the MT growth and shrinkage. Indeed, super coarse-grained Brownian Dynamics modeling benchmarked against *in vitro* experiments measuring MT-based pulling and pushing forces points to such a possibility [27]. Nevertheless, accurate energetic insights are currently missing.

The non-equilibrium nature of the MT tip relaxation in our simulations and improved sampling compared to previous studies provides additional time-resolved information to address this question. In particular, one would expect that, if MTs grow when lateral contacts are strong and disassemble when they are weakened, lateral contacts exposed at the GDP-MT tip would disengage first, followed by PF twist-bending, while they would stay more intact for the GTP-MT tip. To test this idea, we calculated both total MT Solvent Accessible Surface Area (SASA) and total lateral contact area over the course of our simulations (Fig. 3*A*; see Methods). In accordance with the final structures in Fig. 1*C*, GTP- and GDP-MTs gain ~1.4% and ~4.0% of their initial SASA due to splaying during the first 1 *μ*s, respectively. Unexpectedly, however, lateral contact splaying contributes only ~59% and ~20% to the total gain, respectively. This finding implies that MTs loose ~21% of their lateral contacts during the first 1 *μ*s, regardless of the nucleotide state. In addition, we compared the characteristic relaxation time of the MT SASA and the total contact area (Fig. 3*A*; see Methods), which revealed that the former is almost an order of magnitude faster for both GTP- and GDP-MTs. It is therefore mainly the PF twist-bending that promotes the increase of the MT SASA, which in turn causes the lateral contacts to dissociate (at least on microsecond time scales). Interestingly, despite the stronger tendency of GDP-PFs to twist-bend, we still do not observe significantly less lateral contacts relative to the initial contact area at the end of the GDP-MT simulations as compared to the GTP-MT simulations.

**Figure 3:**
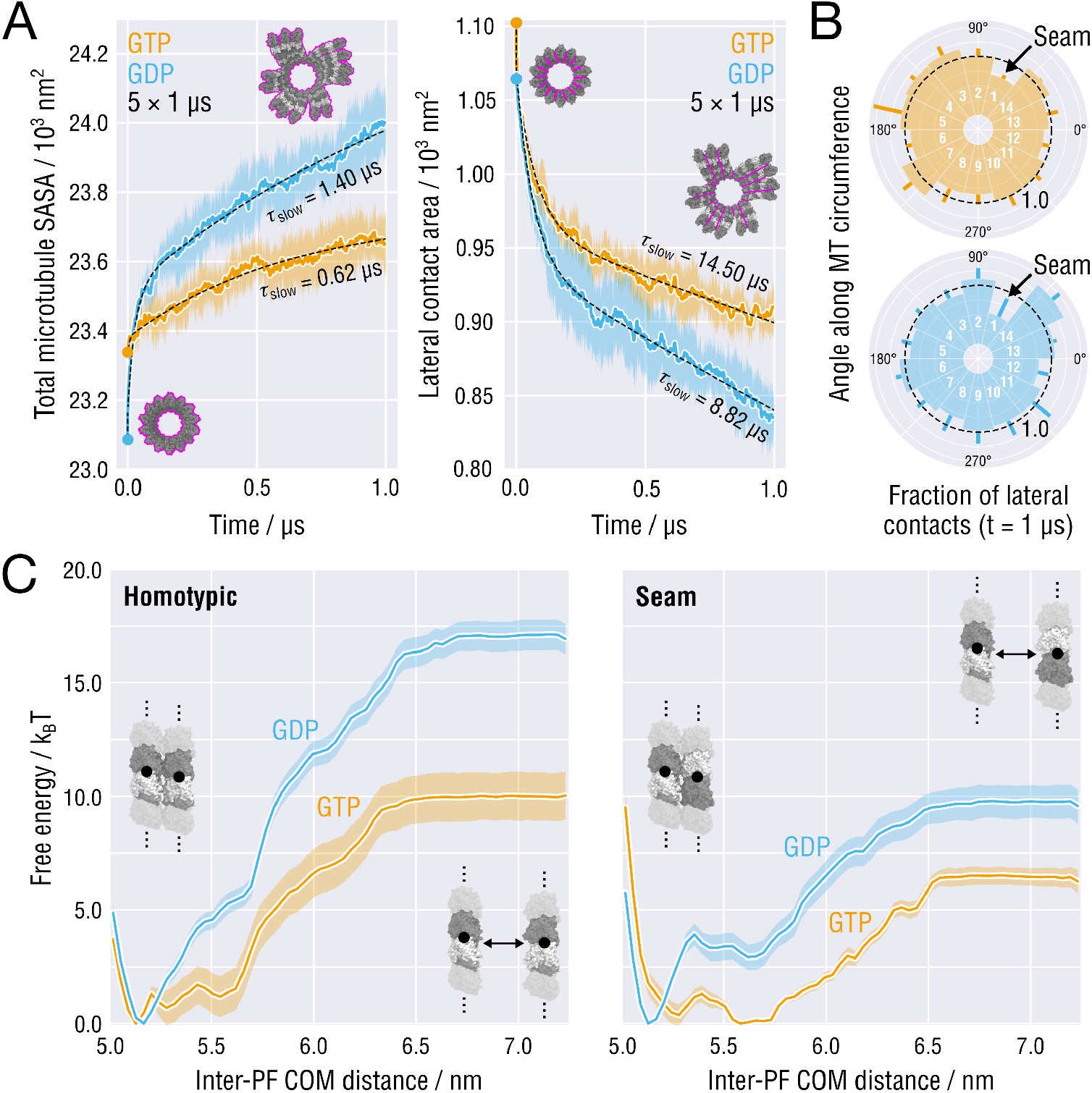
Time dynamics, heterogeneity and energetics of MT tip splaying. **(A)** Time evolution of the total MT Solvent Accessible Surface Area (SASA; left) and the lateral contact area (right), both averaged over the five independent tip relaxation runs shown in Fig. 1*C* (same color coding for GTP- and GDP-MTs was used). Black dashed lines depict fits with a two-component exponential decay function. Characteristic decay times of the slow components are indicated as inserts. **(B)** Lateral contacts preserved after 1 *μ*s of simulation relative to the initial state (indicated with a black dashed line). Positions of the seam are indicated as inserts. **(C)** Free energy profiles of homotypic (left) and seam-like (right) tubulin-tubulin lateral interactions in straight infinitely long PFs as a function of the inter-PF center of mass (COM) distance. Color coding as in **(A)** and **(B)**. Statistical errors are shown as shaded areas (see Methods).

The MT geometry imposes that at one specific lateral interface, called the *seam, α*- and *β*-tubulins form heterotypic contacts (Fig. 1*B*). Structural studies have speculated that the seam is a weak interaction site in the MT lattice and may represent evolutionary tuning to reduce the intrinsic stability of the regular MT lattice [56,54,52]. Indeed, introducing additional seams into the dynamic MT lattice substantially increases the catastrophe frequency and the depolymerization rate, yet not the growth rate [56]. To determine the contribution of the seam to the overall loss of lateral interfaces, we calculated the fraction of pairwise PF contacts preserved after 1 *μ*s of simulation (Fig. 3*B*; see Methods). Interestingly, some lateral contacts – despite the global trend – increased their interaction surfaces during relaxation. We think this is caused by the uneven distribution of PF twist-bending strains at the tip, which transiently pushes the PFs into each other. Regardless of the nucleotide state and in all relaxation runs, the seam PFs disengaged significantly faster than the average of the homotypic PFs (a preserved fraction of 0.43-0.59 compared to ~0.80 for the average homotypic interface), supporting the view that MT plus-ends have a higher tendency to splay at the seam. It is however unlikely that the seam is the primary initiator of lattice opening, as claimed originally, because statistically, it is still more favorable to crack open at any of the more numerous homotypic interfaces.

It seems counterintuitive that the relative loss of lateral interfaces per microsecond is almost equal between the two nucleotide states (Fig. 3*A,B*), while the GDP-MT exerts a much stronger tearing load on the lateral contacts due to the higher PF bending-torsional stiffness (Fig. 1*B,C,D* and Fig. 2*A*). Moreover, it is puzzling why, then, the MTs splay faster at the seam while the seam PFs presumably have bending-torsional properties not largely different from the rest of the MT shaft. Because the preparation of and the simulation conditions for the GTP-MT and GDP-MT models were identical, these observations might point to substantial differences in the free energy landscapes of lateral PF-PF interactions, namely, stronger lateral interactions between GDP-PFs compared to those between GTP-PFs and stronger homotypic lateral interactions compared to seam-like ones. To test this hypothesis, we calculated the lateral interaction free energy between straight PFs locked in the MT shaft as a function of nucleotide state and contact topology. To this end, we used umbrella sampling with initial configurations obtained from 1-*μ*s unrestrained MD simulations of the straight double-PF system described previously [57,35] (see Methods). Initial configurations of the homotypic and seam PFs were derived from the full MT lattice model (Fig. 1*B*). In the umbrella simulations, an external force applied to the inter-PF center of mass (COM) distance, mimicking the effect of PF splaying at the tip, was used to restrain the PFs at a desired distance from one another. Convergence analysis served to ensure that the sampling was sufficient to statistically discriminate between the four tested conditions (Fig. S6).

The resulting profiles shown in Fig. 3*C* reveal that initially – for all tested cases – the free energy rises steeply as the PFs move away from each other from their equilibrium distance ~5.2 nm to roughly 6.5 nm. Clearly, this free energy barrier stabilizes the lateral contact around its natural conformation and explains why sufficient external load applied by the PFs curling outwards is needed for the MT tip to crack open and reach a flared topology. After ~6.5 nm the free energy levels off, which is likely indicative of a high-energy transition state in which the straight PFs have disengaged completely, and a further reduction of the free energy would only be possible through PF twist-bending, which is forbidden by construction in the chosen double-PF setup.

More importantly, as can also be seen in Fig. 3*C*, the energy required to cause the PFs to disengage depends to strongly on both nucleotide occupancy of the *β*-tubulin subunits and topology (seam vs. homotypic). For the homotypic lateral contact, the system in the GDP state has to reach a substantially higher energy Δ*G*\* ≈ 17.1 k_B_T prior to rupture compared to only Δ*G*\* ≈ 10.0 k_B_T for the system in the GTP state. This result becomes particularly interesting in the context of our MT relaxation simulations: although the GDP-MT tip exerts higher tearing loads on its lateral contacts, the load is compensated by the higher lateral stability, which may lead to apparently similar relaxation time scales of the lateral contact fraction for GTP- and GDP-MTs, yet not the total MT area (Fig. 3*A,B*). Even though our data does not directly explain the origin of the observed 7-k_B_T difference in the lateral contact stability, we do not think this energy difference results from altered surface charge distributions or solvation shells because of the high structural similarity of the pre- and post-hydrolysis states of PFs (both initially differ by only ~0.2-nm compaction in the longitudinal dimension). Rather, it is conceivable that – like in the case of longitudinal contacts (Fig. 2*B*) – changes at the inter-dimer interfaces cause subtle structural changes in the vicinity of lateral contacts, hence raising the lateral contact free energy.

For the seam lateral contact, the relationship between the barrier height and the nucleotide state changes drastically (Fig. 3C, right panel). The free energy of the transition state depends only weakly on the nucleotide state (ΔΔG* ≈ 3.3 k_B_T) and is also systematically lower than that of the homotypic system. Thus, given the same tearing load, a seam interface likely would rupture faster than a homotypic interface, which is in qualitative agreement with the SASA analysis (Fig. 3*B*). Also here the observed contact weakening can be most likely ascribed to the non-canonical contact structure, because the seam PFs themselves are not expected to have largely different mechanical properties.

## Conclusions

From non-equilibrium atomistic simulations of complete MT plus-end tips, we have derived a thermodynamic and kinetic description of the MT plus-end tip splaying dynamics, providing a mechanistic interpretation for recent structural experiments. The simulated splayed shapes of MT tips (Fig. 1*C,D*) are generally consistent with cryoelectron tomography measurements [25,26,17], further supporting the view that blunt straight MT tips are strongly energetically unfavored, irrespective of the nucleotide state. Interestingly, cryoelectron tomography shows that PFs bend almost entirely within a radial plain containing the MT axis (out-of-plane deflection of <4 nm over 30-50 nm of PF length; [25]), whereas our simulated structures show somewhat larger PF deflections (Fig. 1*D*; ≲10 nm and ≲15 nm over 40-50 nm of PF length for GTP- and GDP-MTs, respectively). We believe the larger deflections are due to the fact that most of the PFs in our non-equilibrium simulations are transiently stabilized in a metastable state by lateral interactions, unlike those captured recently by cryoelectron tomography [25,27]. Such metastable states with ‘sticky’ protofilament clusters are, however, directly seen in some tomographic reconstructions of both GMPCPP-stabilized and dynamic MT tips [24,26]. Residing in a metastable state with partially coupled PFs during initial lattice splaying might transiently produce an uneven distribution of PF twist-bending and tangential swing strains at the tip and cause a global clockwise twist of the entire MT tip Fig. 1*C*). Twisted MT tips with partially associated PFs are possibly transient, functionally relevant, and nucleotide-dependent intermediates, the exact role of which for the large-scale and long-term MT dynamics is yet to be elucidated. To the best of our knowledge, this behavior has not been accounted for and/or predicted by any of the published physical models of MT dynamic instability.

Our conformational analysis shows that substantially more free energy is required to spontaneously form straight GDP-PFs than straight GTP-PFs at the MT tip (Fig. 2*A*), in good agreement with the recent experimental observation that subpopulations of straight GDP-oligomers occur extremely rarely [58] as well as with recent computational studies [31,32,33,35,36]. Moreover, not only does GTP hydrolysis appear to affect the confrontational rigidity of PFs, but its influence is also highly anisotropic, with GTP-PFs being more flexible with respect to both outward twist-bending and tangential swing and GDP-PFs being more rigid, but particularly rigid to motions in the tangential direction. We hypothesize that the pre- and post-hydrolysis modes of PF motion have different effects on the collective dynamics of the MT tip. During MT assembly, flexible GTP-PFs have an elevated rate of both spontaneous straightenings and encounters with neighboring GTP-PFs, while the higher twist-bending and, especially, tangential rigidity of GDP-PFs reduces this rate and causes high tearing loads on lateral interfaces during MT disassembly.

The observed dependence of the mechanical properties of individual PFs on the nucleotide state may be a prototypical example for an inter-subunit cooperativity that does not involve significant backbone rearrangements, but is capable of causing large-scale conformational effects, as recently demonstrated for other protein nanomachines such as the F_1_-ATPase [59] or the bacterial MreB filaments [60]. We find that, also in the MT system, this communication is mediated by clusters of strong pairwise residue interactions concentrated mainly near the nucleotide binding pocket as well as the H8 helix, the H8-H7 loop, and the accompanying H11’ helix at the intra- and inter-dimer interfaces (Fig. 2*B,C* and Fig. S3).

The free energy calculations in conjunction with the non-equilibrium relaxation simulations of complete MT tips enable us to address a key question about the role of lateral contacts in the MT plus-end stability. While the presented estimates of the lateral interaction energies are magnitude-wise consistent with previous ones derived from atomistic simulations and/or enhanced sampling [61,34], we have been able to discern in detail the effects of the nucleotide state on the homotypic lateral contact stability owing to ~100-fold more MD sampling (Fig. 3*C*, left). We have found that GTP hydrolysis, in fact, increases the lateral contact stability by ~7 k_B_T, and contrary to our initial skepticism, the subsequent convergence analysis has confirmed that (Fig. S6). Moreover, the fact that both GTP- and GDP-MT tips loose approximately the same fraction of lateral contacts within 1 *μ*s of simulation (Fig. 3*A*, right panel) – despite the much higher conformational strain stored initially in the GDP-MT tips – can only be explained by more stable lateral contacts between GDP-PFs. Though counterintuitive, we believe that the hydrolysis-driven stabilization of lateral contacts in the MT lattice, accompanied by the increase of longitudinal strain in PFs, is a necessary condition for regulating MT dynamics and triggering MT catastrophe for the reasons explained below.

Timing the proper sequence of tubulin binding, PF straightening, lateral contact formation, and GTP hydrolysis at the MT plus-end tip is essential for progressive MT growth, but also for the onset of MT catastrophe. The rates of both spontaneous PF straightening and lateral association between PFs (≲10^5^ s^−1^ [62,63,61,34,27,35]) are, however, substantially larger than the rates of tubulin turnover and GTP hydrolysis at the MT tip (0.24 – 40 s^−1^ at 10-15 *μ*M free tubulin [64,16,17,18]). This implies that PF curvature fluctuations and PF-PF interactions are unlikely to limit MT growth and shrinkage. Thus, the mechanism of MT catastrophe is perhaps more complex than it was proposed by early allosteric models postulating that GTP-tubulin binding causes straightening of PFs required for the assembly of MTs and GTP hydrolysis induces curvature in PFs inconsistent with the MT lattice [65,66], or by more recent proposals that the lateral contacts weakened by GTP hydrolysis can no longer counteract the strain energy stored in the MT lattice [49,37,55,39].

Instead, given the thermodynamics of the plus-end PFs and the kinetics of MT lattice splaying (Figs. 1–3), the stability of the MT tip structure might be determined by the fast dynamic equilibrium between the straight(er) coupled and curved uncoupled states of neighboring PFs (schematically illustrated in Fig. 4*A*). We propose that the tendency of each PF to twist-bend and disconnect from its nearest neighbors would be balanced out by the lateral interactions, resulting in a complex potential energy landscape of the MT tip having multiple minima separated by energy barriers. One of these minima would correspond to fully splayed MT tips akin to the ones observed experimentally [25,25,27], while the many others to tips with metastable clusters of less curved PFs (Fig. 1*C*). Both minima depths and barrier heights would be tightly controlled by the nucleotide, resembling the concept of information-driven Brownian motors or ‘ratchets’ [67], where the ‘information’ is the dependence of the rates of PF straightening and PF-PF contact formation on the nucleotide state and the directed motion (MT growth) is achieved by rectification of thermal noise. Therefore, if building PF clusters was favorable and the barriers for their formation were low (*e.g*., at a GTP-MT tip), the MT tip would then favor, by mass action, the pathway through these metastable states towards the straight MT lattice, provided there is sufficient and timely delivery of GTP-tubulin to the tip. Otherwise, if the barriers were high and PF cluster formation unfavorable (*e.g*., at a GDP-MT tip), the net MT growth would no longer be possible even under sufficient GTP-tubulin concentrations, resulting in a catastrophe event.

**Figure 4:**
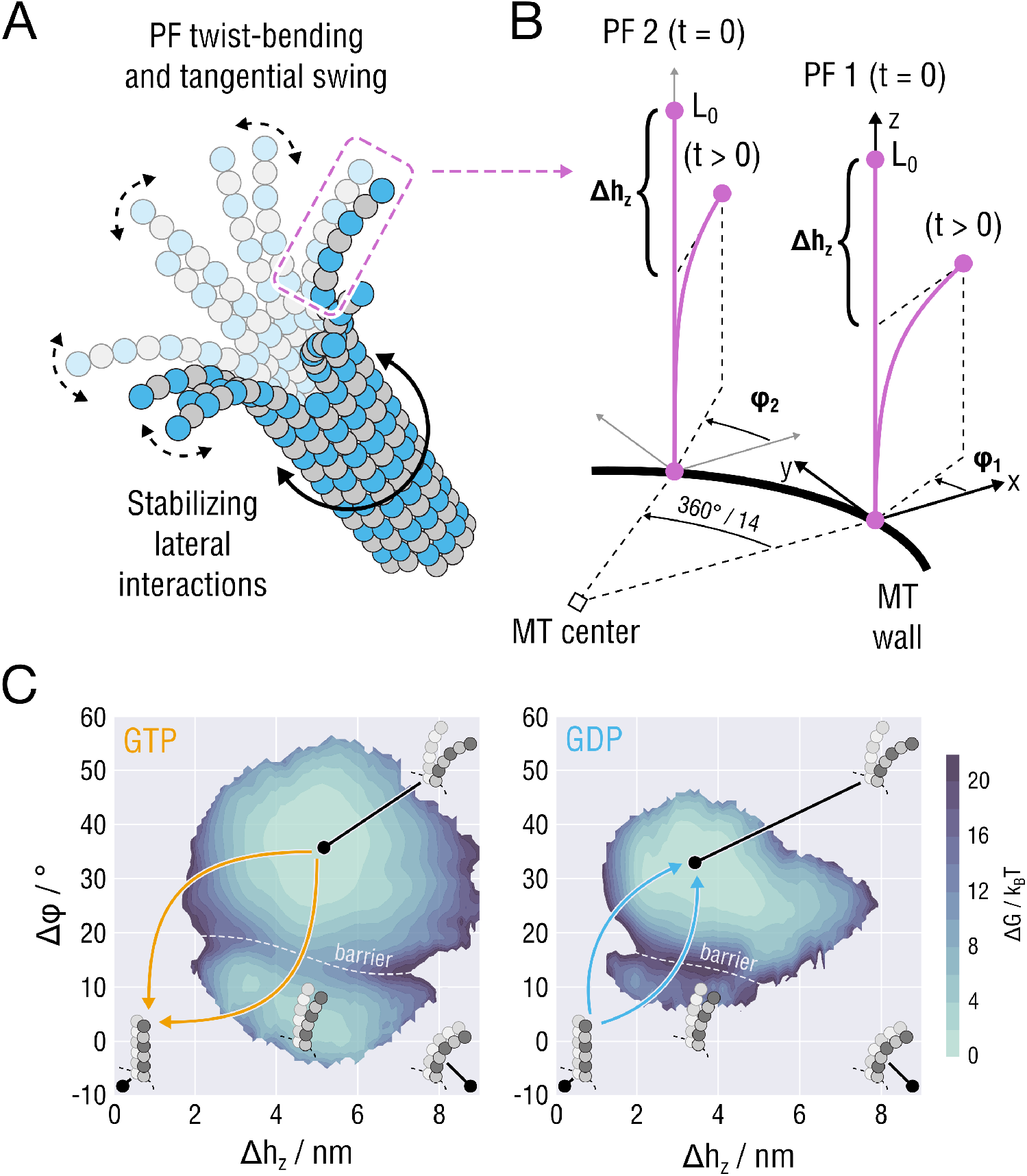
Schematic illustrating the proposed mechanism of the MT plus-end stability. **(A)** Exemplary structure of the MT tip determined by the competition between the stabilizing inter-PF interactions and the twist-bending dynamics of individual PFs. **(B)** Schematic visualization of a pair of neighboring PFs (purple lines). For illustration, other PFs at the MT tip are not considered and the MT body, to which the PFs are rigidly attached, is shown as a black thick line. Both the distance between the PFs and their length are drawn not-to-scale for clarity. Given this simplified representation, the order parameters that determine the extent of PF outward bending and splaying are the PF height difference, *Δh_z_*, and the difference in the azimuthal angles, Δ*ϕ* = *ϕ*_2_ – *ϕ*_1_. **(C)** Free energy profiles describing the bending and splaying dynamics of the double-PF system as a function of the nucleotide state, generated by resampling the results of Fig. 2*A* and Fig. 3*C* (see Methods).

To illustrate how this mechanism works, we consider a simplified subsystem consisting of only two neighboring PFs at the MT tip (Fig. 4*A,B*). In principle, the knowledge of the energetics of PF twist-bending and lateral PF-PF association can be merged in order to provide a joint free energy landscape of the coupled PF dynamics (the reconstruction procedure is described in Methods). Briefly, our procedure defines the PFs conformations by two order parameters: the PF height difference, Δ*h_z_*, reflecting the extent of twist-bending and the azimuthal angle difference, Δ*ϕ* = *ϕ*_2_ – *ϕ*_1_, reflecting the extent of tangential splaying. Figure 4*C* shows the reconstructed free energy landscapes for the motion of the coupled PF system in each nucleotide state. First, as expected, both landscapes exhibit two free energy minima reflecting the states of fully uncoupled and splayed PFs and partially coupled straightened PFs. Second, according to the landscapes, two distinct pathways for the formation of the straight MT wall would be feasible: (a) partial straightening of both PFs through partial lateral association, *i.e*. first crossing the barrier and then spontaneously forming a straight lattice from the metastable state; or (b) simultaneous and full straightening of both PFs accompanied by full lateral association. In the scenario (b), no metastable PF cluster would be formed, however, at the price of a much higher free energy required. Finally, GTP hydrolysis modulates the shape of the free energy landscape such as to make the formation of both metastable clusters and straight MT wall more unfavorable (Fig. 4*C*, left *vs*. right panel). Specifically, the energy barrier that the system would have to overcome to achieve the straight MT state raises considerably owing to both higher GDP-PF rigidity and higher PF-PF interaction free energy. These simplified considerations corroborate that the inability of the GDP-MT tip to elongate might be primarily caused by the strongly suppressed collective rate of GDP-PF straightening and lateral contact formation at the tip (‘backward’ transition), rather than by the intrinsic tendency of the GDP-MT tip to disassemble (‘forward’ transition), hence the description as a ‘ratchet’.

Finally, the role of the seam on MT dynamics has been elusive. Here we have shown that, although MT tips crack open at different lateral interfaces, the rate of PF splaying at the seam is significantly elevated (Fig. 3*B*), in accordance with the original hypothesis by Nogales and colleagues [54,52]. This observation is further supported by the calculated lateral free energies (Fig. 3*C*, right panel) that only weakly depend on the nucleotide state and are systematically lower than those for the homotypic interface. These results confirm that the functional effect of the seam might be exactly the opposite of a homotypic interface: namely, to provide a kinetic destabilization of the entire MT tip by lowering the energy barrier for lateral contact dissociation, especially during MT shrinkage (Fig. 4*C*, right panel). This mechanism would explain why MTs containing more than one seam exhibit more frequent catastrophe events and shrink faster than single-seam dynamic MTs [56], yet the excess presence or absence of seams does not necessarily affect the bending rigidity of MTs [68]. Rather, the presence or absence of weaker seams may offer an additional mechanism to regulate MT polymerization and depolymerization kinetics.

## Methods

### Construction of MT tip models

Initial models for the tubulin dimers were obtained from PDB IDs 3JAT (GMPCPP) and 3JAS (GDP) [52] by extracting the central dimer from the 3 × 2 lattice patches (chains A and H in the original PDBs). GMPCPP was converted into GTP by replacing the carbon atom between *α*- and *β*-phosphate with an oxygen atom. The missing loop in the *α*-subunit (residues 38-46) was modelled in for structure consistency using MODELLER version 9.17 [69] but excluded from further refinement. Similarly to our previous study [35], the flexible C-termini (*α*:437-451 and *β*:426-445) were not included in our simulations to reduce the system size and to achieve the best possible sampling given the system size.

Asymmetric (C1) reconstructions of 14-PF GMPCPP- and GDP-MTs decorated with kinesin (kindly provided by R. Zhang; [52]) were used to construct the all-atom models of MT lattices. Although the C1 reconstructions have a lower resolution (4.0-4.1 Å compared to 3.5 Å for the symmetrized ones), they reflect the correct seam location and topology as well as the slightly increased PF-PF distance at the seam. Subsection of the cryo-EM maps enclosing two layers of dimers in the axial direction were extracted, and copies of the atomistic dimer models were rigid-body fitted into the subsection densities (Fig. S4). The constructed models were solvated in a triclinic water box of size 33.2 × 33.2 × 25.8 nm^3^ and subsequently neutralized with 150 mM KCl.

Refinement was done with correlation-driven molecular dynamics implemented as a custom module in the GROMACS 5.0.7 package [70,41], following our previously published protocols [41,35]. Briefly, we employed a cold-fitting protocol (NPT ensemble, T = 100 K, p = 1 atm) containing three phases: short restraint-free equilibration for 10 ns, refinement with a resolution and force constant ramp for 35 ns, and finally, 15 ns of simulated annealing. The sigma parameter controlling the simulated map resolution was ramped from 0.6 nm to 0.2 nm. The force constant was ramped from 1 × 10^5^ kJ/mol to 1 × 10^6^ kJ/mol.

The final MT plus-end models were constructed by grafting three copies of the refined subsection models onto one another by applying a translation operation to each copy that conforms to the dimer periodicity (also known as *dimer rise*; ~8.31 nm for GTP-MTs and ~8.15 nm for GDP-MTs [52,40]) and removing the terminal *α*-tubulin monomers from the plus-end (see Fig. 1*B*). The resulting tip structures (79 *α*-tubulins and 84 *β*-tubulins; ~56 nm in length) were kept blunt to maximize the initial lateral contact area.

### Relaxation simulations of MT tips

The CHARMM22* force field [71] and the CHARMM-modified TIP3P water model [72] were used in all simulations. GTP/GDP parameters were adapted from those for ATP/ADP implemented in the CHARMM22/CMAP force field [72,73]. Titration curves of histidines were calculated using the GMCT package [74] and assigned as described previously [31].

The final MT tip structures were centered and resolvated in a larger periodic box of size 50 × 50 × 63 nm^3^ to accommodate larger splayed conformations and to avoid collisions of the PFs belonging to periodic images (at least on microsecond time scales). As in the refinement simulations, the solvated tip structures were neutralized with 150 mM KCl, yielding an atomic system with ~15.6 million atoms. All subsequent MD simulations were carried out with GROMACS 2019 [75]. Lennard-Jones and short-range electrostatic interactions were calculated with a 0.95-nm cutoff, while long-range electrostatic interactions were treated using particle-mesh Ewald summation [76] with a 0.12-nm grid spacing. The bond lengths were constrained using the LINCS algorithm [77] (hydrogen bonds during initial equilibration and all bonds in the production runs). Velocity rescaling [78] with a heat bath coupling constant of 0.5 ps was used to control the temperature for solute and solvent separately. Applying virtual site constraints [79] allowed us to increase the integration step size to 4 fs in the production runs. Center-of-mass correction was applied to solute and solvent separately every 100 steps. To mimic the minus-end attachment, we applied position restraints with the force constant *k* = 350 kJ/mol/nm^2^ to the bottom layer of monomers (9 *α*-tubulins and 5 *β*-tubulins in total).

With the above parameters fixed, the equilibration protocol consisted of the following steps: (i) energy minimization using steepest descent; (ii) short NVT equilibration for 1 ns at T = 100 K with position restraints on all heavy atoms and using a 1-fs integration time step; (iii) gradually heating up the system to 300 K within 5 ns in the NVT ensemble using a 2-fs integration time step; (iv) equilibration in the NPT ensemble for 5 ns (Berendsen barostat [80] with a 5-ps coupling constant) using a 2-fs time step; (v) gradual release of the position restraints (except those mimicking the minus-end attachment) within 10 ns in the NPT ensemble (isotropic Parrinello-Rahman barostat [81] with a 5-ps coupling constant) using a 4-fs time step; (vi) pre-equilibration for 5 ns in the NPT ensemble (same as in (v)) using a 4-fs time step. The last frame of step (vi) was used to spawn 5 independent 1-*μ*s relaxation runs for each nucleotide state.

### Simulations of single finite PFs and infinite coupled PFs

Straight GTP-PFs and GDP-PFs comprising three longitudinally coupled tubulin dimers were extracted from a seam-distant location of the respective MT tip models (Fig. S5). The single-PF systems were then solvated in a water box of size 13.5 × 13.5 × 28.0 nm^3^ and neutralized with 150 mM KCl. The box size was sufficient to avoid collisions with periodic images. The PFs were then equilibrated as described above and the equilibrated structures were used to span 20 independent 4-*μ*s production runs for each nucleotide state.

Double-PF systems were prepared as described in our recent study [35]. Briefly, for each nucleotide state, we extracted two 2 × 2 tubulin patches, one from a seam-distant location mimicking a homotypic interface and the other from the seam (Fig. S5). The patches were then solvated in a water box of size 12.7 × 12.7 × 21.0 nm^3^ and neutralized with 150 mM KCl. These finite double-PF models were converted into ‘infinite’ double-PF models by removing the extra tubulin dimers and nucleotides. Water and ion atoms were then trimmed to conform to the experimental value of the axial periodic dimension, namely, ~8.31 nm for GMPCPP-MTs and ~8.15 nm for GDP-MTs [52]. The number of ions in the trimmed water shell was fixed such as to keep the systems neutral and to maintain the ionic strength of 150 mM KCl. The double-PF systems were equilibrated as described above with the only difference that we used the semi-isotropic Parrinello-Rahman barostat to control the system size fluctuations in the axial (along the PF) and transverse (perpendicular to the PF) directions separately. The equilibrated structures were used to span a 1-*μ*s production run for each nucleotide state.

### Calculation of the PF-PF association free energy profiles

Our approach to calculate the lateral PF-PF interaction free energies relied on a previous study [34]. Clearly, the 1-*μ*s production runs were not sufficiently long to yield the full interaction profiles because high free energies would be required for the PFs to disengage. Therefore, we employed the umbrella sampling technique [82] in conjunction with the Weighted Histogram Analysis Method (WHAM) [83,84]. We first defined the inter-PF COM distance to be the reaction coordinate. The biasing potential was tuned to be 4000 kJ/mol/nm^2^ and was applied to restrain only the transverse component of the inter-PF distance. To cover the full range of inter-PF interactions, the distance between 5.0 nm and 7.2 nm was split into windows, each being separated by 0.05 nm from its nearest neighbors. This partitioning of the reaction coordinate space yielded sufficient overlap between neighboring windows in the absolute majority of cases. In those cases, where the overlap was still ≲30%, new windows were added. The production runs were used for ‘seeding’ the umbrella simulations, where each ‘seed’ was separated from all the others by at least 50 ns in time. Seeding structures for those windows that were not initially covered by the unrestrained simulations of the double-PF systems were derived from neighboring windows located 0.05 nm away in the reaction coordinate space. Each window was then simulated for 1 *μ*s and the autocorrelation times of the inter-PF COM fluctuations were assessed. If an autocorrelation time was above the average, the respective umbrella window simulation was extended to 1.3 *μ*s. For determining the convergence of the free energy profile for each condition (nucleotide state and contact topology), we calculated the profiles using WHAM as a function of the length of umbrella window trajectories with an increment of 100 ns (Fig. S6). This analysis confirmed that the final free energy profiles shown in Fig. 3*C* are sufficiently converged (maximum deviation from the previous increment <1 k_B_T). Free energy uncertainties were estimated using Bayesian bootstrapping [84] of the complete histograms scaled by inefficiency factors *g_i_* = 1/(1 +*τ_i_*), where *τ_i_* is the autocorrelation time of umbrella window *i* [85,86].

### Calculation of PF traces

PF traces were calculated to analyze the deflections of individual PFs in a radial plain passing through the MT central axis and an azimuthal plane orthogonal to the MT central axis. To this end, a discrete contour was drawn on each PF structure using specific reference points, here referred to as *nodes*. The definition of the nodes depended on the type of the longitudinal contact (*i.e*., intra-dimer or inter-dimer) as well as on the position of the tubulin dimer in the PF (i.e., terminal or non-terminal). For each nonterminal intra-dimer interface, the node was defined as the COM of the residues *α*:405-411 and *β*:249-264. For each non-terminal inter-dimer interface, the COM of *α*:251-266 and *β*:395-401 was used. A schematic structure of a PF trace defined using this methodology is shown in Fig. S7. The residues *α*:251-266 and *β*:395-401 were used to define the minus- and plus-end nodes, respectively.

### Principal component analysis of PF dynamics

To quantify the highest-variance (lowest-energy) uncorrelated motions of the single PF conformational dynamics, the principal component analysis [87] was carried out. First, for each independent single PF trajectory (see above), we extracted all backbone atoms, while excluding the first 1 *μ*s of the respective trajectory from the analysis. Second, all backbone trajectories were superimposed relative to the minus-end *α*-tubulin. Then, all independent trajectories belonging to the same nucleotide state were concatenated and the atomic displacement covariance matrix was calculated. The covariance matrix was diagonalized to yield the eigenvalues {λ_*i*_} and eigenvectors {**v**_*i*_}. Finally, the concatenated trajectory for each nucleotide state was projected onto the first and second eigenvector of the this matrix. The projections were then histogrammized to obtain the equilibrium distributions *p*(*ξ*_1_,*ξ*_2_) converted to the free energy profiles *G*(*ξ*_1_, *ξ*_2_) = – k_B_T ln *p*(*ξ*_1_, *ξ*_2_) (Fig. 2*A*). The extent of conformational flexibility of GTP-GDP-PFs was estimated by summing up the respective eigenvalues 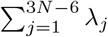, where *N* is the number of backbone atoms. The relative contribution of eigenmode *i* can then be expressed as 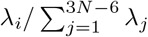.

### Joint free energy landscape of coupled PF dynamics obtained through resampling

To assess the combined effect of PF elasticity and PF-PF interaction on the dynamics of neighboring PFs, we employed a resampling scheme using the data available from the single-PF simulations (Fig. 2*A*) and the umbrella sampling of lateral interactions (Fig. 3*C*). To mimic the presence of a lateral neighbor, we first copied the full ensemble of PF conformations and rotated the copy by 2*π*/14 radians along the MT circumference away from the original (Fig. 4*B*). We then introduced a simplified set of parameters to describe the conformations of the PF neighbors: namely, the PF height difference, Δ*h_z_* = *L*_0_ – *h_z_*, where *L*_0_ is the straight PF height, and the PF splaying angle, Δ*ϕ* = *ϕ*_2_ – *ϕ*_1_. The parameter Δ*h_z_* reflects the difference between the heights of the straight and a curved PF conformation, *i.e*. the extent of PF twist-bending. The height of each PF is defined as the position of the COM of the plus-end *β*-tubulin along the MT main axis. In turn, the splaying angle Δ*ϕ* is the difference between the azimuthal angles *ϕ*_1_ and *ϕ*_2_ defined by projecting the positions of the COMs of the plus-end *β*-tubulins onto the transverse plane. The angle *ϕ*_2_ is measured in the coordinate system of the first PF such that *ϕ*_2_ – *ϕ*_1_ is almost always positive.

With the order parameters Δ*h_z_* and Δ*ϕ* defined, the resampling procedure was as follows. The parametric space (Δ*h_z_*, Δ*ϕ*) was partitioned into bins of finite size. For each bin 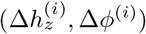, all corresponding PF conformations were found and extracted from the PF ensembles (original and copy), and all possible pairs of PFs were constructed without replacement. For each PF pair *k* in bin *i*, the lateral interaction free energy was computed as 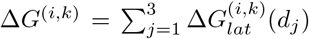, where {*d_j_*} are the COM-COM distances between the neighboring dimers and Δ*G_lat_*(*d*) is the lateral interaction free energy landscape (Fig. 3*C*, left panel). The weighted probability distribution of the PF pair conformations that also accounts for lateral interactions between the PFs was then obtained by bin counting, where each observation *k* in bin *i* was weighted by *w*^(*i,k*)^ ∝ exp[– Δ*G*^(*i,k*)^/k_B_T]. The weighting ensures that those PF conformations showing a strong lateral interaction contribute more to the total probability distribution, thus creating a metastable state at low Δ*ϕ* values.

### Force distribution analysis of longitudinal intra- and inter-dimer interactions

The force distribution analysis was performed using a modified version of GROMACS 2020.4 that includes an implementation of the FDA module v2.10.2 [50,51] (https://github.com/HITS-MBM/gromacs-fda). For the visual analysis of force distributions (Fig. S3), forces between pairs of residues – located at intra- and inter-dimer interfaces – were calculated and averaged over the concatenated PF trajectories for each nucleotide state. A cutoff of 50 pN was applied to the absolute force values to highlight only strong clusters. For the purpose of analysis, solvent and KCl atoms as well as atoms involved in virtual site constraints were removed from the concatenated trajectories. Post-hydrolysis changes in the force distributions (Fig. 2*B*) were calculated by subtracting the average GTP distribution from the average GDP distribution and filtering out changes smaller than 50 pN and larger than −50 pN. These changes were then mapped onto the PF structures and color coded according to the sign of the changes.

### Calculation of RMSD, total MT SASA and lateral contact area

The MT relaxation trajectories were stored every 125000 steps (500 ps). These trajectories were analyzed for the extend of conformational changes such as RMSD, total MT SASA, lateral contact area. The time-resolved RMSD values were calculated relative to the starting straight configuration shown in Fig. 1*B* using a GROMACS internal tool [88]. The time-resolved total MT SASA, *SASA_MT_* (*t*), was calculated in a memory-efficient way using a GROMACS internal tool [89]. The lateral contact area was expressed as a sum over the SASA of all PFs separately minus the total MT SASA divided by two, that is, 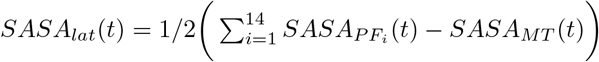.

## Supporting information

Atomic model of the GTP-MT tip

Atomic model of the GDP-MT tip

GTP-MT tip (run 1, top view)

GTP-MT tip (run 2, top view)

GTP-MT tip (run 3, top view)

GTP-MT tip (run 4, top view)

GTP-MT tip (run 5, top view)

GTP-MT tip (run 1, side view)

GTP-MT tip (run 2, side view)

GTP-MT tip (run 3, side view)

GTP-MT tip (run 4, side view)

GTP-MT tip (run 5, side view)

GDP-MT tip (run 1, top view)

GDP-MT tip (run 2, top view)

GDP-MT tip (run 3, top view)

GDP-MT tip (run 4, top view)

GDP-MT tip (run 5, top view)

GDP-MT tip (run 1, side view)

GDP-MT tip (run 2, side view)

GDP-MT tip (run 3, side view)

GDP-MT tip (run 4, side view)

GDP-MT tip (run 5, side view)

Twist-bending mode (side view)

Twist-bending mode (top view)

Tangential swing (side view)

Tangential swing (top view)

## Data analysis and availability

All post-processing calculations were done using GROMACS internal tools, Python 2.7 [90], and Numpy [91]. Graphs were produced with the Matplotlib v2.2.5 [92] and Seaborn v0.9 [93] libraries. All structure and map manipulations were performed using UCSF Chimera v1.14 [94] or VMD v1.9.3 [95]. The VMD software was further used for visualization of all MT and PF structures as well as for producing the movies. Atomic coordinates of the initial MT tip models are provided as Supplementary Files 1 and 2. The raw MD trajectories (> 1 TB of data) that support the findings of this study are available from the corresponding authors upon request.

## Acknowledgements

We acknowledge financial support from the Max Planck Society (M.I. and H.G.) and the German Research Foundation via the grant IG 109/1-1 (awarded to M.I.). Computational resources were provided by the Max Planck Computing and Data Facility and the Leibniz Supercomputing Centre (Garching, Germany).

