## Supplementary Material for "Bending-torsional elasticity and energetics of the plus-end microtubule tip"

[1]Theoretical and Computational Biophysics,
Max-Planck-Institute for Biophysical Chemistry,
Am Fassberg 11, D-37073 Göttingen, Germany

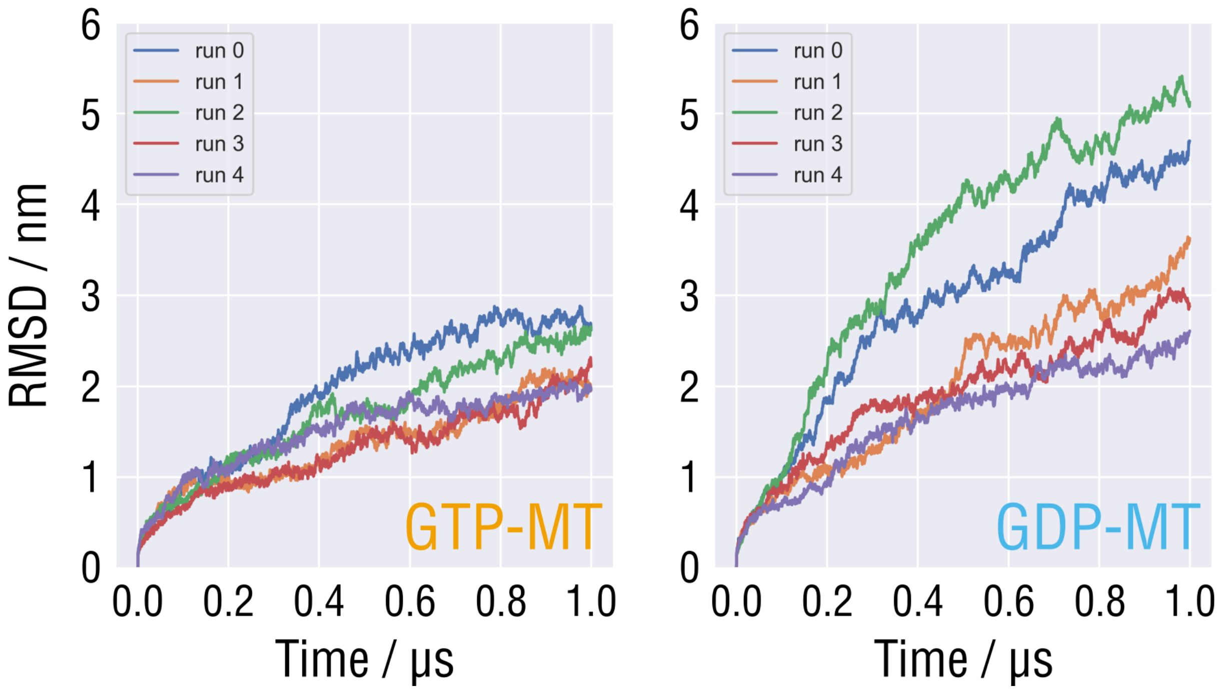


**Fig. S1.** Root mean square deviation (RMSD) of the relaxing MT tip structure from the initial straight conformation for each independent run and in each nucleotide state.

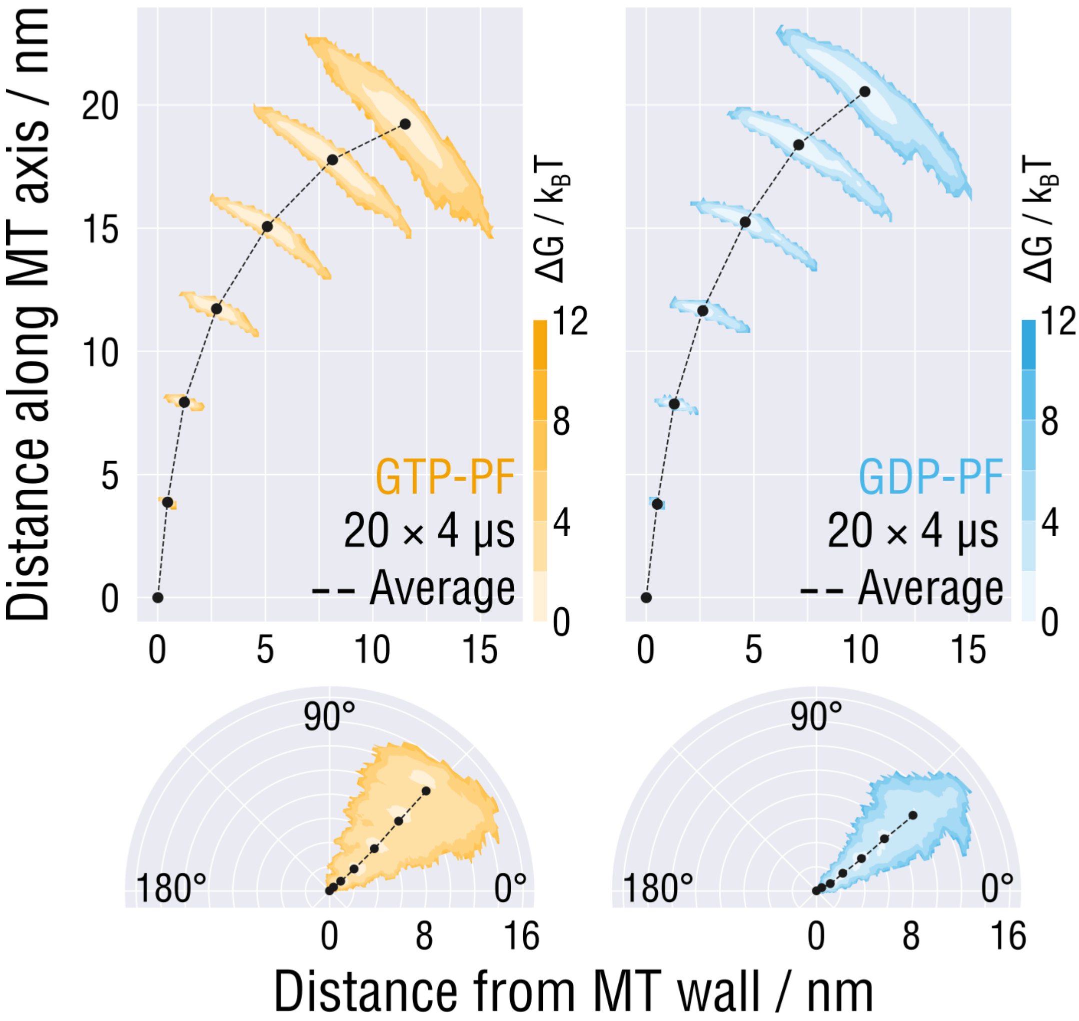


**Fig. S2.** Equilibrium distributions of PF traces aligned with respect to the first minus-end monomers and projected onto the radial (top row) and transversal (bottom row) planes.

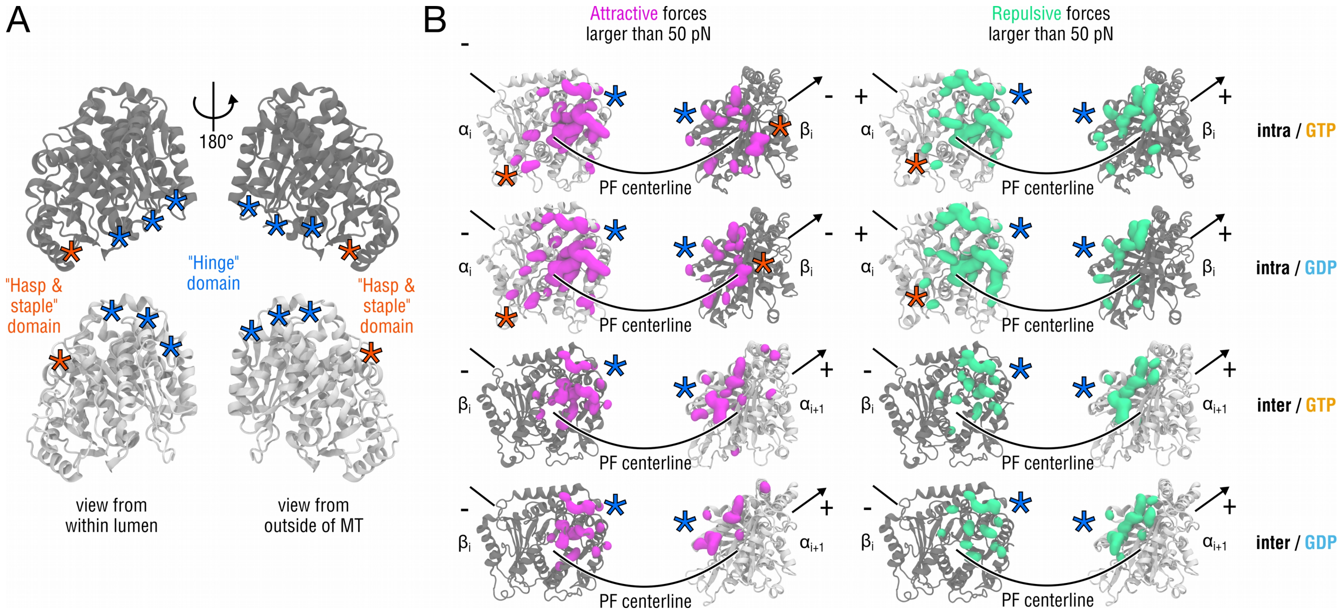


**Fig. S3. (A)** Schematic showing the location of key intra- and inter-dimer interaction domains termed 'hinges' (marked with red stars) and 'hasp & staple' (marked with blue stars). **(B)** Intra-dimer (two top rows) and inter-dimer (two bottom rows) contact clusters of pairwise interaction forces obtained by the Force Distribution Analysis (FDA). Amino acids contributing to attractive interactions (negative forces) are marked with magenta blobs, whereas those contributing to repulsive interactions (positive forces) are marked with green blobs. A lower limit cutoff of 50 pN for both negative and positive pairwise forces was applied to highlight strong interaction clusters. α- and β-tubulins are shown as silver and gray ribbons, respectively.

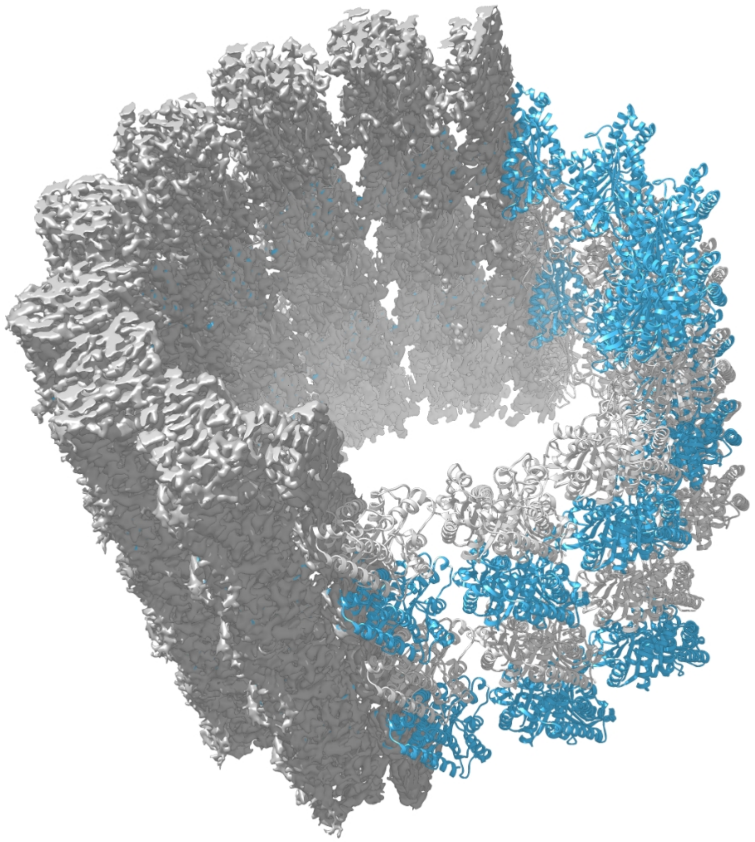

**Fig. S4.** Shown is the cross-section of the cryo-EM map section used in the refinement simulation filled with rigid-body fitted copies of the atomistic dimer model.

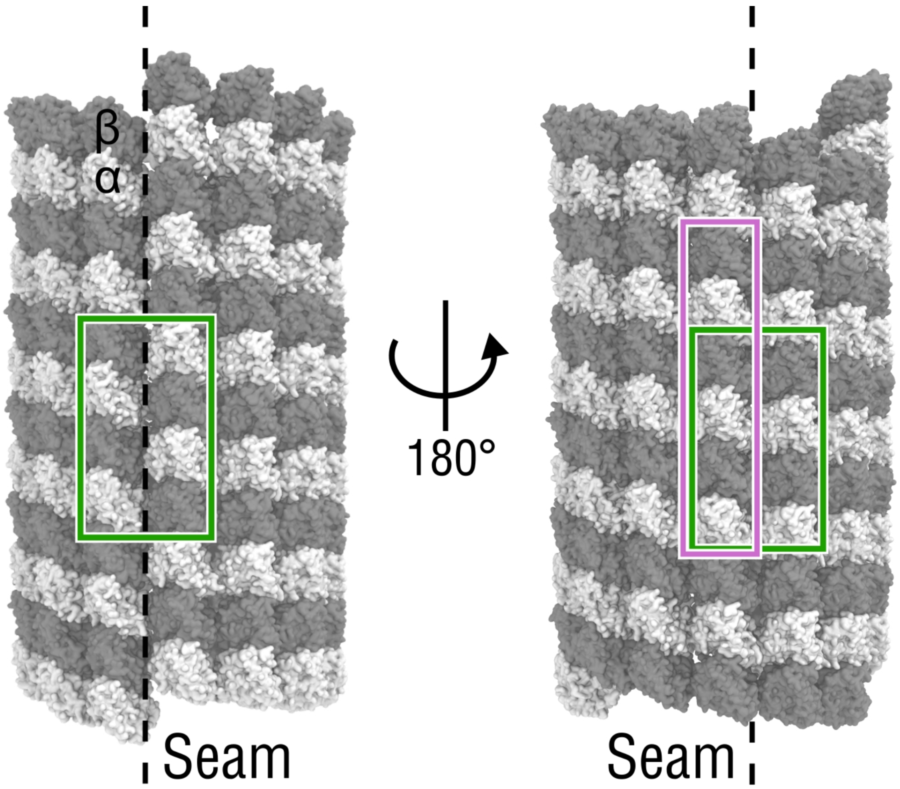


**Fig. S5.** Locations of the part of the full MT tip model used to extract the starting structure for the single-PF simulations (purple rectangle) as well as of the seam and seam-distant parts used to extract the 2×2 tubulin patches for the umbrella sampling simulations of lateral PF-PF interactions (green rectangles).

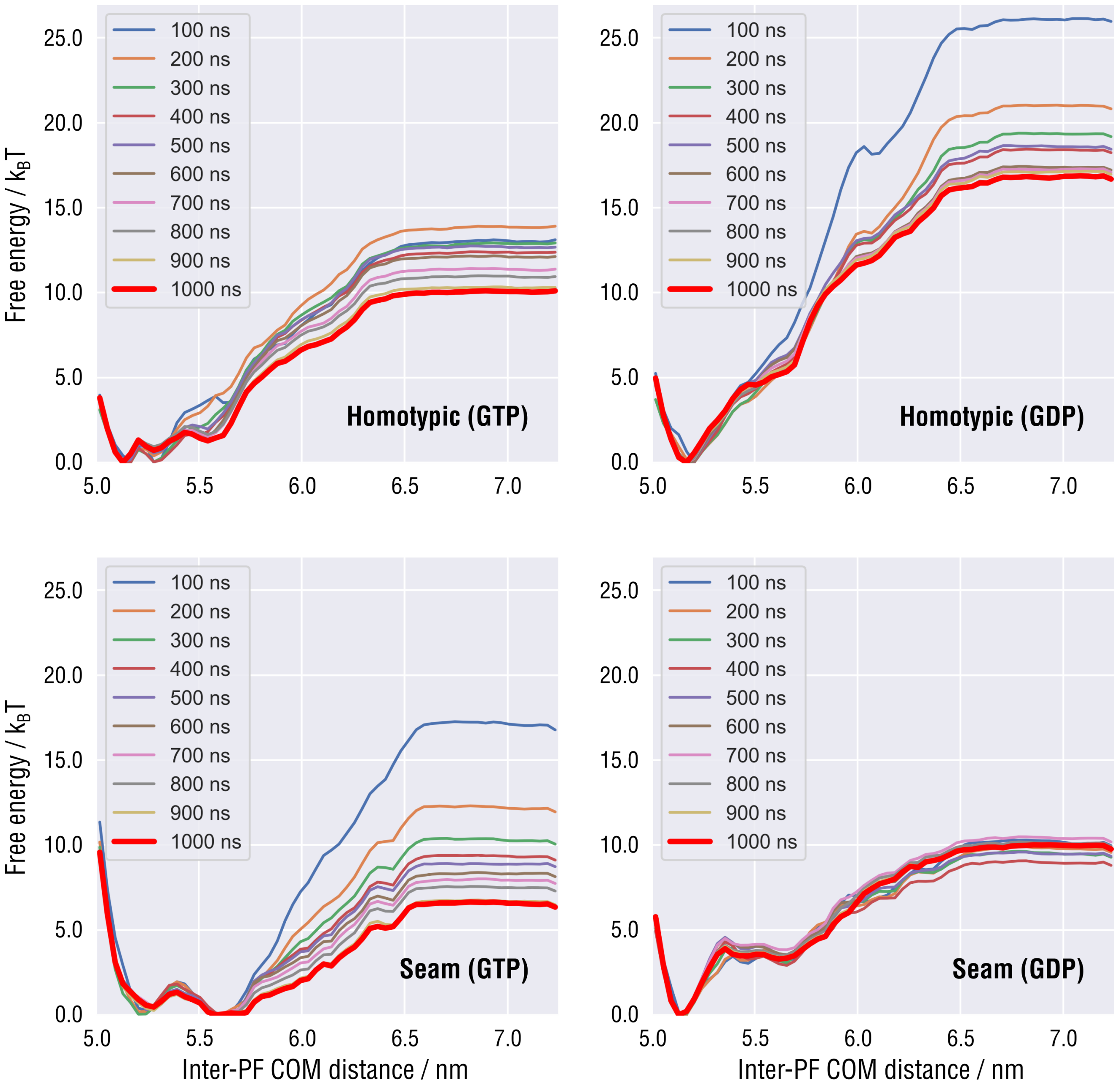


**Fig. S6.** Convergence of the lateral interaction free energy landscapes for both nucleotide states and contact topologies with the length of umbrella sampling trajectories taken for the analysis. Except for the one corresponding to the seam contact in GDP state (bottom right), all of the free energy landscapes initially overestimate the later interaction free energy and then gradually converge to the steady-state profiles as the amount of sampling increases. For the final profiles (red thick lines), the maximum deviation from the previous increment (900 ns per window) is less than 1 $k_B T$.

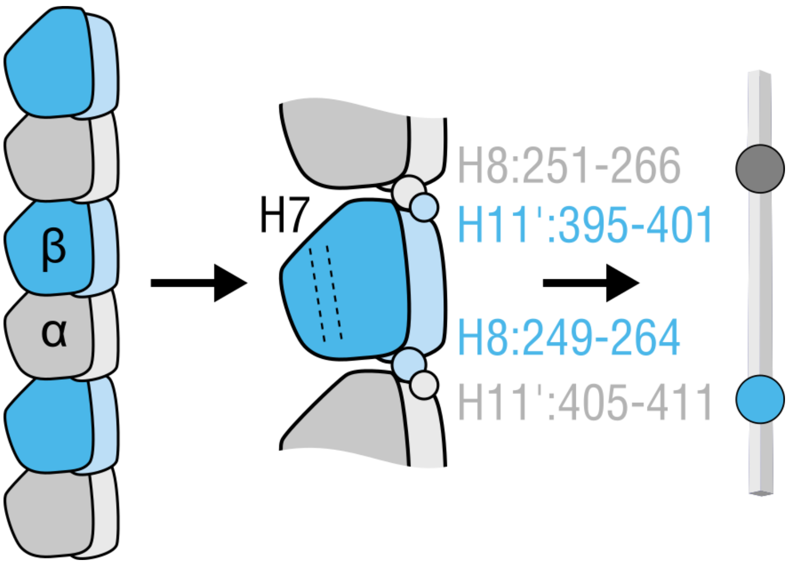


**Fig. S7.** Schematic illustrating the definition of a PF trace.

**Supplementary Movies S1-S20.** Visualizations of the relaxation process of the simulated GTP- and GDP-MT tips. For each independent simulation, a top view (from the plus-end) and a side view (facing the seam) were recorded.

**Supplementary Movies S21-S24.** Visualizations of the twist-bending and tangential 'swing' modes of PF motion. For each mode, a top view and a side view were recorded.

**Supplementary Files 1 and 2.** Atomic coordinates of the initial GTP- and GDP-MT tip models in the PDB format.
